# pbcftools: parallel execution of bcftools for large variant call sets

**DOI:** 10.64898/2026.08.03.742604

**Authors:** Ge Zhang

## Abstract

**Summary:** bcftools is the standard toolkit for handling VCF and BCF variant files, but it processes records on a single core; its --threads option speeds up only compression of the output, not the work done on variant records. Processing large call sets is therefore slow, and users often divide the genome and reassemble the results by hand. We present pbcftools, a Perl wrapper that does this automatically: it splits the genome into chunks, runs an ordinary bcftools command on each in parallel, and reassembles the outputs by a method suited to the data type. Across Linux servers, Windows/WSL2 workstations and Apple laptops, with bcftools 1.21 to 1.24, parallel output was identical to serial output for every command tested. On 1000 Genomes Phase 3 data, operations writing compressed VCF ran 10.8 to 21.1 times faster with 32 cores and up to 35.4 times with 64, those writing text 3.7 to 12.8 times, and merging 100 VCF files 19.2 times. pbcftools also runs on LSF and Slurm clusters.

**Availability and implementation:** pbcftools is written in Perl (>= 5.16) and requires bcftools; local parallel execution also requires Perl module Parallel::ForkManager. It is released under the MIT license at https://github.com/zhangge-uc/pbcftools (DOI: 10.5281/zenodo.21780361).

## 1 Introduction

Bcftools (Danecek *et al*. 2021) and the underlying HTSlib library (Bonfield *et al*. 2021) are the standard tools for handling variant call data sets. Their record-processing work, including filtering, normalization, annotation, summary statistics and merging, runs on a single core. The --threads option speeds up only BGZF compression of the output stream, but not the parsing and computation applied to each variant record. As call sets have grown to tens of millions of sites and thousands of samples, this has become the main reason routine steps are slow.

Most bcftools operations treat each genomic position independently, which makes them *embarrassingly parallel*: applying a command separately to non-overlapping chunks of the genome and joining the results gives the same answer as processing the whole file in serial. Users already exploit this manually, such as using GNU parallel (Tange 2015), shell loops or workflow managers. However, they must then generate the regions usually by chromosomes, launch one command per region, track the output of each command, preserve record order, and choose a correct way to combine the results. Each step differs by command, and mistakes are easy to make and hard to notice. Distributed frameworks such as CloudMerge (Sun *et al*. 2018) and workflow managers such as Nextflow (Di Tommaso *et al*. 2017) and Snakemake (Mölder *et al*. 2021) can address parts of this, but the infrastructure or scripting are not accessible to ordinary users.

To solve this problem, we introduce pbcftools, a command line Perl wrapper that performs the whole cycle of bcftools job division, execution and reassembly using a similar interface to bcftools – normal bcftools command line arguments with a few additional options to control parallelization; everything else is passed to bcftools untouched.

## 2 Implementation

pbcftools parallelizes bcftools operations that are embarrassingly parallel, and its scope is bounded by two conditions: *first*, the work must be divisible along the genome, either by interval, when each record is handled independently of its neighbors, or by whole chromosome or contig, when the computation carries state along a sequence but is independent between chromosomes or contigs; *secondly*, the results must be recombinable without further computation, either by concatenation, as for tabular text and for VCF/BCF records, or by a simple summary, as for the counters produced by stats command. Operations failing either condition, such as whole file operations like sort and index, or operations with output files difficult to combine, such as cnv and isec, are passed to bcftools unchanged. Table 1 lists how pbcftools treats each bcftools command.

**Table 1.**
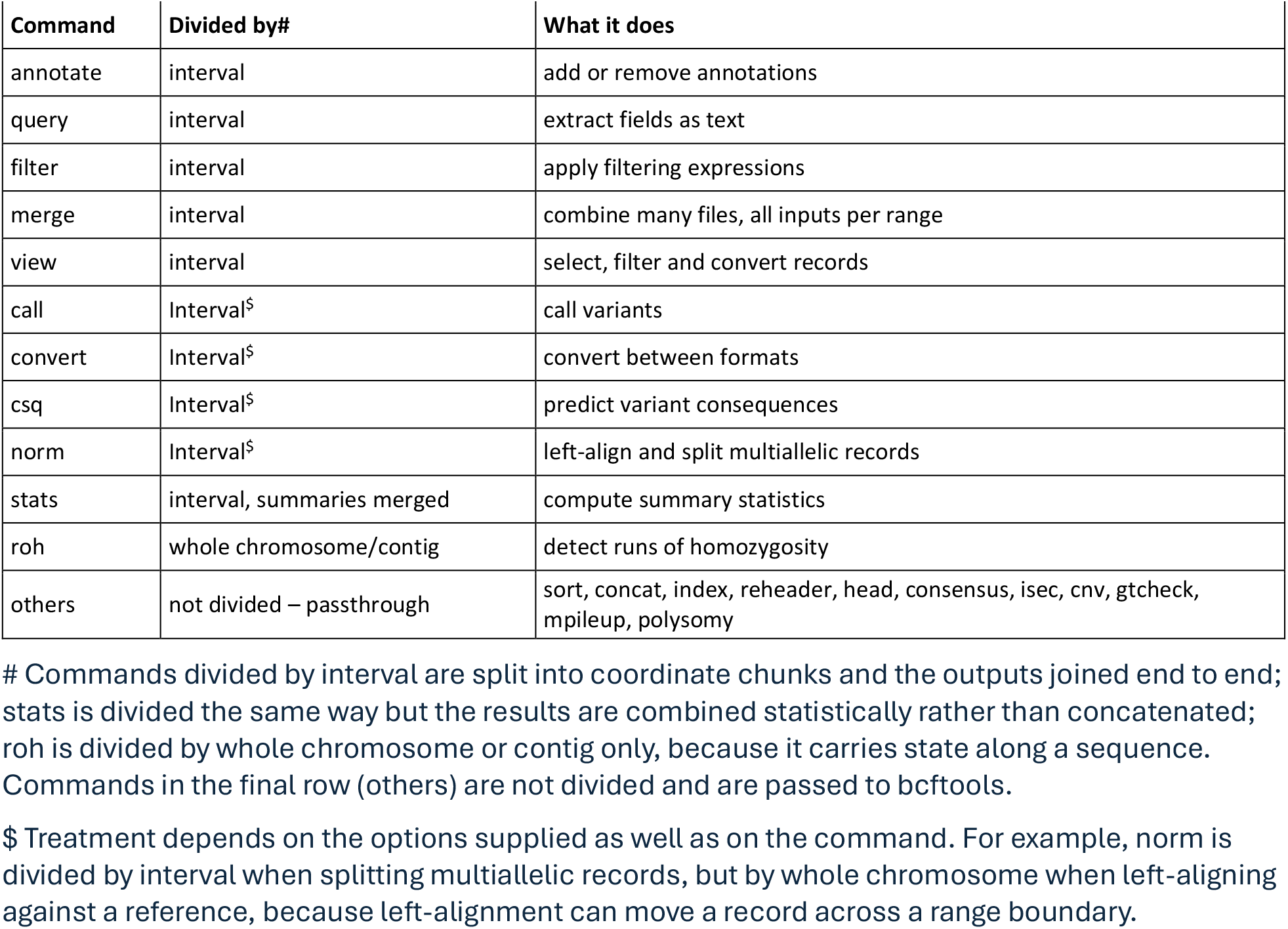
How pbcftools treats each bcftools command.

To ensure familiarity and consistency with bcftools, pbcftools takes normal bcftools command line arguments and its own options, all begin with --p_ for controlling parallelization behavior (such as --p_jobs for number of concurrent jobs). As no bcftools option starts with ‘p_’, the two sets of options cannot collide. Each command is then assigned one of four treatments (Table 1): division by interval, division by interval with summaries merged (stats), division by whole chromosome or contig (roh, which would otherwise report one run of homozygosity as two shortened ones), or ordinary serial execution (cannot be parallelized, passthrough to bcftools).

To split the work by intervals or chromosomes/contig, pbcftools first needs the name and length of every sequence in the input file(s). It takes these from the first source that provides them: the companion index of the input (the .csi or .tbi file), the contig lines of the VCF/BCF header, a user-supplied FASTA index (--p_fai), or a built-in table of human chromosome sizes (--p_ref). The last two pbcftools options are used when the input VCF file(s) carries no contig lengths in their header. The whole genome, or regions specified by the user (by bcftools option -r, --regions or -R, --regions-file FILE), is divided into chunks of a configurable size (--p_len). Each variant is assigned to exactly one chunk, so that joining the pieces reproduces the bcftools result exactly, and the user’s own overlap setting is preserved at the start of the requested region.

The chunks run as separate processes on one machine, or as LSF or Slurm jobs on a HPC cluster, where a sliding window keeps a fixed number of parallel jobs (set by --p_jobs) in flight. They are then combined by a method suited to their content: compressed variant files (VCF or BCF) are concatenated in naïve mode, tabular text is joined with a single header block, and stats summaries are merged with plot-vcfstats. If any chunk fails, pbcftools writes no output and reports an error. Output is assembled in a working directory and moved into designated output place only after every chunk has succeeded, so an interrupted run cannot leave a partial file with silently failed regions.

## 3 Validation

Correctness was assessed at three levels, all reproducible using the testing code in the GitHub repository. A self-contained testing suite builds small synthetic files and runs 22 commands through both bcftools and pbcftools to validate consistency of the results, and conducts extensive checks of failure handling, such as that a failed run leaves an existing output file untouched and that an interrupted run never leaves a partial file. A real-data validation and benchmark were conducted using 1000 Genomes Phase 3 data (The 1000 Genomes Project Consortium *et al*. 2015), including whole-genome sites-only file (84.8M variants) and chromosome 1 genotypes for 2,504 samples (6.5M variants).

Every comparison between the serial bcftools and pbcftools parallel runs with different numbers of concurrent jobs passed on all the machines tested, including Linux servers, Windows/WSL2 workstations and Apple laptops, with bcftools 1.21 to 1.24 and Perl 5.34 to 5.40, together with LSF and Slurm clusters. Results did not depend on the operating system, the processor architecture or the bcftools version; only the achievable speed differed.

The benchmark results (Table 2) showed that the performance gain of pbcftools over bcftools is mainly determined by the amount of computation per record, and it has two sources. The first is writing the output: operations producing compressed VCF reached 10.8 to 21.1-fold at 32 cores, because compression is itself substantial work that divides along with the records. This holds whether or not the input carries samples, so splitting multiallelic records in a sites-only file reached 15.8-fold. The second is per-sample processing: among operations writing text, those reading genotypes reached 12.1 to 12.8-fold, whereas extracting a few fields from a site-only file reached only 3.7-fold, having almost no computation to divide. As shown by Figure 1, the gain scales near linearly with the number of concurrent jobs until the available cores are saturated.

**Table 2.** Speed relative to serial bcftools on a Linux server with 96 cores (Intel Xeon Gold 6248R, Fedora 40, bcftools 1.22)

| Operation (command) | Type<sup>#</sup> | Output<sup>\$</sup> | Serial<sup>%</sup><br>(sec) | 4 cores | 8 cores | 16 cores | 32 cores |
| --- | --- | --- | --- | --- | --- | --- | --- |
| extract genotypes to text file<br>(query -f '%CHROM\t%POS[\t%GT]\n') | genotype | text | 941.2 | 3.22× | 5.77× | 9.64× | 12.20× |
| genotype summary statistics<br>(stats) | genotype | text | 149.6 | 2.24× | 4.17× | 7.58× | 12.77× |
| subset three samples<br>(view -s \$SAMPLES) | genotype | VCF | 528.6 | 3.04× | 5.93× | 11.48× | 21.14× |
| extract fields as text<br>(query -f '%CHROM\t%POS\t%REF\t%ALT\t%INFO/AF\n') | site | text | 104.7 | 1.06× | 1.79× | 2.72× | 3.69× |
| site summary statistics<br>(stats) | site | text | 102.0 | 1.24× | 2.40× | 4.54× | 8.23× |
| filter on allele frequency<br>(filter -i 'INFO/AF>0.05 && INFO/AF<0.95') | site | VCF | 133.3 | 1.61× | 3.14× | 5.94× | 10.75× |
| remove an annotation<br>(annotate -x INFO/AF ) | site | VCF | 277.8 | 1.90× | 3.73× | 7.24× | 13.69× |
| split multiallelics<br>(norm -m -both) | site | VCF | 353.6 | 2.30× | 4.50× | 8.74× | 15.82× |

<sup>#</sup>: Operations labelled genotype were run on chromosome 1 of the 1000 Genomes Phase 3 genotype data (2,504 samples, 6.5M variants) using 1 Mb ranges; those labelled site were run on the whole-genome sites-only file (84.8M variants) using 10 Mb ranges. <sup>\$</sup>: The Output column distinguishes operations that write compressed VCF from those that write a tabular or summary text file, which is the main determinant of how well an operation parallelizes. <sup>%</sup>: Serial times are single runs of bcftools on the same machine measured in seconds. Every comparison produced records identical to the serial result.

**Figure 1.**
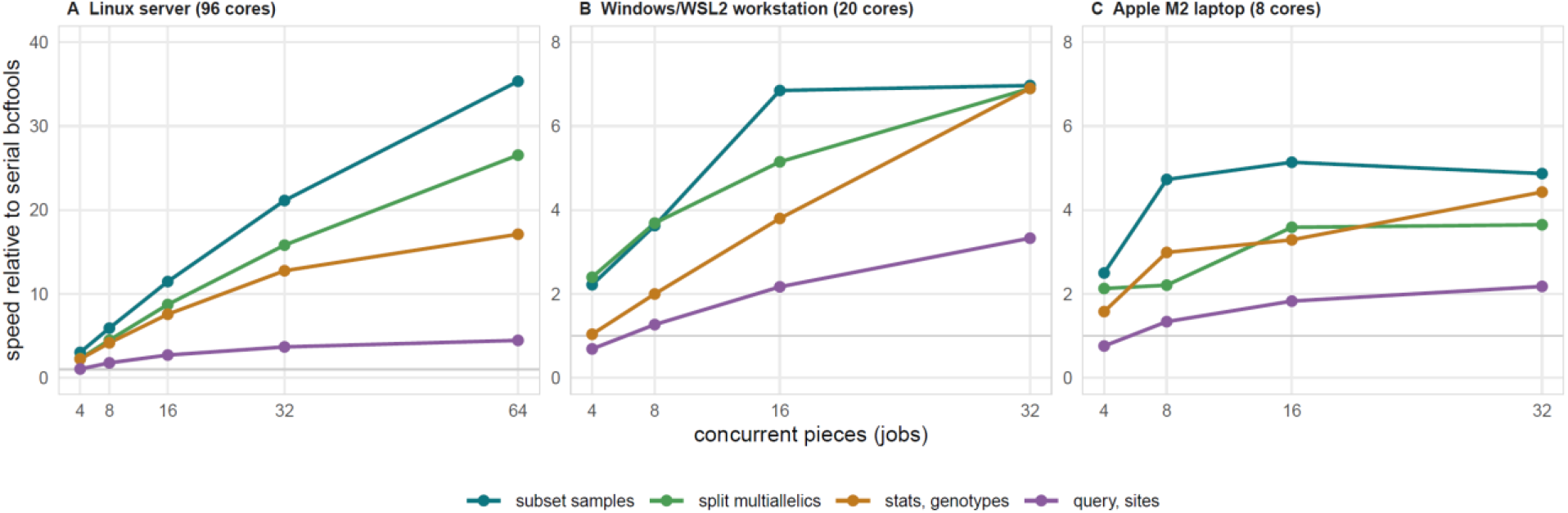
Speed relative to serial bcftools as the number of concurrent pieces increases, for four representative operations. (A) a 96-core Linux server, (B) a 20-core Windows/WSL2 workstation, and (C) an 8-core Apple M2 laptop, where the x axis is the number of concurrent jobs. On the server the gain rises nearly linearly while cores remain available; on the workstation and laptop the curves flatten once concurrency approaches the core count.

Merging benefits substantially from parallelization, because serial bcftools merge must open all inputs and read them together throughout. Merging 100 VCF files, each with ten samples across chromosome 1 of the 1000 Genomes Phase 3 genotype data (6,468,094 variants) on a 96-core linux server using pbcftools is 19.2-fold faster with 64 concurrent jobs than serial run. The gain of performance of the same merging test is 14.8-fold at 256 jobs on a LSF cluster despite the cost of job queueing. Cluster timings should be read differently from single-machine ones, since each piece waits in a queue and that wait is included in the measurement. Cluster execution therefore pays when each job is substantially longer than a typical queue wait.

## 4 Discussion

pbcftools removes the single-core bottleneck in bcftools for the large class of operations that treat genomic positions independently, while producing identical records and preserving the bcftools command line. It complements bcftools --threads, which only parallelizes compression within one process, whereas pbcftools divides the data processing work across processes.

Three points bear on how much benefit to expect. First, the gain follows the computation carried by each record and the size of the job. Operations writing compressed output (e.g. compressed VCF and BCF) and those involving more computation per record gain most (over 20-fold); extracting a few text fields from a site-only VCF file gained less than 4-fold. Second, the size of each range matters for light operations and hardly at all for computationally intensive jobs such as merging. Ranges must be large enough that real work outweighs the cost of starting a process, and numerous enough to occupy every core; for merging, varying the range size twenty-five-fold changed the result by less than 4%.

Third and most importantly, parallel execution by pbcftools moves the bottleneck from the processor to the storage. Using only a single core, bcftools cannot process records as fast as the storage can supply them, so the disk is left waiting. Spreading computational work across many cores removes the limit, and the combined demand of those cores then meets what the storage can deliver. As shown by Figure 1, the performance of most operations increase nearly linearly with the number of concurrent jobs until all the computational cores are occupied – meaning users can comfortably run 48∼64 concurrent jobs if computational cores are available.

Correctness is the property we have worked hardest to make trustworthy, so pbcftools determines each command explicitly for parallelization, refuses commands it does not recognize, and falls back to serial execution (passthrough to bcftools) whenever a command cannot be parallelized or be read unambiguously. Remaining limitations, including the commands run serially by choice and the plugins not yet supported, are documented in the repository.

In summary, pbcftools runs an existing bcftools command in parallel automatically, using the same command line with a few additional options. It is easy to install and runs unchanged on a laptop, a high-performance server or an HPC cluster. On every platform tested, it produced records identical to those of bcftools. By making use of the multiple cores now standard on ordinary machines, it substantially reduces the time required to process large variant call sets.

## Acknowledgements

This work used Anvil at Purdue University through allocation BIO260210 from the Advanced Cyberinfrastructure Coordination Ecosystem: Services & Support (ACCESS) program, which is supported by U.S. National Science Foundation grants #2138259, #2138286, #2138307, #2137603 and #2138296.

## Author contributions

GZ conceived and designed the software, wrote the code, performed the validation and benchmarking, and wrote the manuscript.

## Use of AI tools

Generative AI tools (Anthropic Claude, via the Claude Code command-line interface) were used during software development, testing and manuscript preparation. Their use included code revision and refactoring, development of the test harness, analysis of benchmark results, and drafting and editing of the manuscript text. All software design decisions, validation and benchmark protocols, results and scientific conclusions were determined and verified by the author, who takes full responsibility for the content of this article.

## Availability and requirements

- Project name: pbcftools
- Project home: https://github.com/zhangge-uc/pbcftools (DOI: 10.5281/zenodo.21780361)
- Operating systems: Linux, macOS, Windows (via WSL2)
- Programming language: Perl ≥ 5.16 (tested on 5.34–5.40)
- Requirements: bcftools (tested on 1.21–1.24); plot-vcfstats for parallel stats; Parallel::ForkManager for local parallel execution; a scheduler client and a shared filesystem for cluster modes
- License: MIT

## References

Bonfield JK, Marshall J, Danecek P et al. HTSlib: C library for reading/writing high-throughput sequencing data. GigaScience 2021;10(2):giab007. 10.1093/gigascience/giab007.

Danecek P, Bonfield JK, Liddle J et al. Twelve years of SAMtools and BCFtools. GigaScience 2021;10(2):giab008. 10.1093/gigascience/giab008.

Di Tommaso P, Chatzou M, Floden EW et al. Nextflow enables reproducible computational workflows. Nat Biotechnol 2017;35(4):316–9. 10.1038/nbt.3820.

Mölder F, Jablonski KP, Letcher B et al. Sustainable data analysis with Snakemake. F1000Res 2021;10:33. 10.12688/f1000research.29032.2.

Sun X, Gao J, Jin P et al. Optimized distributed systems achieve significant performance improvement on sorted merging of massive VCF files. GigaScience 2018;7(6):giy052. 10.1093/gigascience/giy052.

Tange O. GNU Parallel 20150322 (‘Hellwig’). Zenodo, 22 Mar. 2015. 10.5281/ZENODO.16303.

The 1000 Genomes Project Consortium, Auton A et al. A global reference for human genetic variation. Nature 2015;526(7571):68–74. 10.1038/nature15393.

